# The properties of buried ion pairs are governed by the propensity of proteins to reorganize

**DOI:** 10.1101/2020.02.03.932012

**Authors:** Christos M. Kougentakis, Lauren Skerritt, Ananya Majumdar, Jamie L. Schlessman, Bertrand García-Moreno E.

## Abstract

Charges are incompatible with the hydrophobic interior of proteins, yet proteins use buried charges, often in pairs or networks, to drive energy transduction processes, catalysis, pH-sensing, and ion transport. The structural adaptations necessary to accommodate interacting charges in the protein interior are not well understood. According to continuum electrostatic calculations, the Coulomb interaction between two buried charges cannot offset the highly unfavorable penalty of dehydrating two charges. This was investigated experimentally with two variants of staphylococcal nuclease (SNase) with Glu:Lys or Lys:Glu pairs introduce at internal i, i+4 positions on an α-helix. Contrary to expectations from previous theoretical and experimental studies, the proteins tolerated the charged ion pairs in both orientations. Crystal structures and NMR spectroscopy studies showed that in both variants, side chains or backbone are reorganized. This leads to the exposure of at least one of the two buried groups to water. Comparison of these ion pairs with a highly stable buried ion pair in SNase shows that the location and the amplitude of structural reorganization can vary dramatically between ion pairs buried in the same general region of the protein. The propensity of the protein to populate alternative conformation states in which internal charges can contact water appears to be the factor that governs the magnitude of electrostatic effects in hydrophobic environments. The net effect of structural reorganization is to weaken the Coulomb interactions between charge pairs; however, the reorganized protein no longer has to pay the energetic penalty for burying charges. These results provide the framework necessary to understand the interplay between the dehydration of charges, Coulomb interactions and protein reorganization that tunes the functional properties of proteins.

## Introduction

Buried ionizable residues in proteins are rare but essential for biochemical energy transduction. When buried alone, the p*K*_a_ values of these residues are shifted in the direction that favors the neutral state, relative to their values in water, consistent with the unfavorable transfer of an ionizable moiety from water to the hydrophobic interior of a protein^1^. In many natural systems, such as in enzyme active sites^2,3^, proton pumps^4,5^, ion channels^6^, and pH-sensing motifs in signaling proteins^7,8^, ionizable residues can be buried in pairs or in networks. Despite being able to form ion pairs^9,10^, continuum electrostatics theory predicts that a favorable Coulomb interaction is not strong enough to compensate for the unfavorable dehydration of two charges^11,12^. Detailed structural and thermodynamic characterization of the energetics of buried ion pairs in natural systems is rarely feasible since many of these proteins are difficult to work with experimentally. The lack of experimental data has hampered the development of computational methods capable of accurately capturing the interplay between favorable Coulomb interactions and unfavorable dehydration energies governing the energetics of ion pair burial. A detailed understanding of charge-charge interactions in hydrophobic environments is necessary to gain mechanistic insight into biochemical energy transduction processes, and to design novel proteins (e.g. enzymes, H^+^ pumps) with these important functional properties.

Recent studies have shown that it is possible to introduce two ionizable residues in the hydrophobic core of the model protein staphylococcal nuclease (SNase) as a charged pair^12^, without significantly affecting the structure of the background protein. When buried alone, the p*K*_a_ values of the introduced Glu-23 and Lys-36 are highly anomalous and favor the neutral state, relative to their values in water. When introduced together, subtle reorganization of backbone and side chain dipoles along with water penetration creates a highly polar environment that stabilizes the charged E23/K36 pair, with a coupling energy of 5 kcal/mol. Previous theoretical studies predicted that the arrangement of dipoles that would stabilize the arrangement of one ion pair would destabilize the reverse orientation^13^. Consistent with this prediction, reversing the introduced ion pair in SNase (V23K/L36E) leads to a protein of marginal stability with a weak coupling energy, as the protein appears incapable of reorganizing dipoles or introducing internal water molecules to stabilize the reverse pair^14^.

In this study, we have characterized the structural and thermodynamic properties of two variants of SNase with internal Lys:Glu or Glu:Lys pairs at positions 62 and 66 on helix-1 of SNase. These positions were chosen because the residues are close in sequence and poised to make stabilizing i,i+4 interactions in both orientations^15,16,17,18,19,20,21^. The p*K*_a_ values of the buried residues in the single variants are shifted to favor the neutral state, relative to their values in water^1,22^. The p*K*_a_ values of the introduced Glu/Lys residues in both T62K/V66E (KE) and T62E/V66K (EK) variants are normalized relative to those measured in the single variants (i.e. more similar to their values in water). These proteins exhibit more complex conformational responses than observed in the 23:36 ion pair variants, which allows the protein to tolerate the burial of charged EK and KE pairs regardless of orientation. The structural response of these proteins demonstrates that charged groups are incompatible with the protein interior even in the presence of stabilizing interactions, and that correctly predicting alternative conformations accessible in the conformational ensemble is necessary to understand how proteins are able to tolerate the burial of charged pairs and networks.

## Results

### Circular dichroism (CD) spectroscopy

Circular dichroism spectra of the EK and KE variants collected at pH values between 5-9 (Figure 1) were indicative of well-folded proteins. The spectra of the KE variant are pH independent and were superimposable with those of the background protein. Conversely the EK variant displays small but reproducible shifts at 208 and 222 nm relative to the background protein across the range of pH values studied, consistent with minor structural reorganization. Similar shifts in the CD spectra are observed in both the T62E and V66K single variants at pH values where their respective introduced residues are ionized (Figure S1).

**Figure 1.**
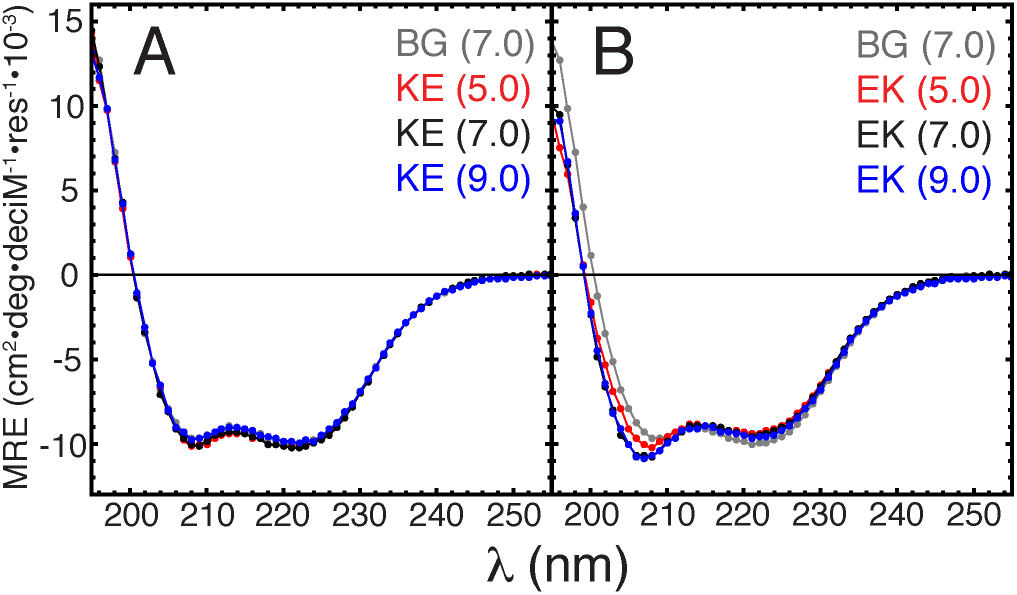
Circular dichroism spectra of background protein, (**A**) KE and (**B**) EK variants as a function of pH. The pH value of each sample is shown in parenthesis.

### Thermodynamic stability

The thermodynamic stabilities of the two ion pair variants were measured as a function of pH using chemical denaturation with guanidinium hydrochloride monitored with Trp. fluorescence (Fig. 2, Table 1 and S1-3). Introduction of an ionizable residue with a perturbed p*K*_a_ leads to a characteristic pH dependence to its thermodynamic stability^1^. Thermodynamic linkage analysis of the difference in the pH-dependence of the thermodynamic stability between the background protein and the variant protein with buried ionizable residues can be used to determine apparent p*K*_a_ values of these introduced residues. Previous studies have shown that buried Lys residues at positions 62 and 66 titrate with p*K*_a_ values of 8.1 and 5.7, respectively, far from the p*K*_a_ value of 10.4 of Lys in water^22^. Buried Glu residues at the same positions have p*K*_a_ values of 7.7 and 8.5, respectively, far from the p*K*_a_ value of 4.5 of Glu in water^1^.

**Figure 2.**
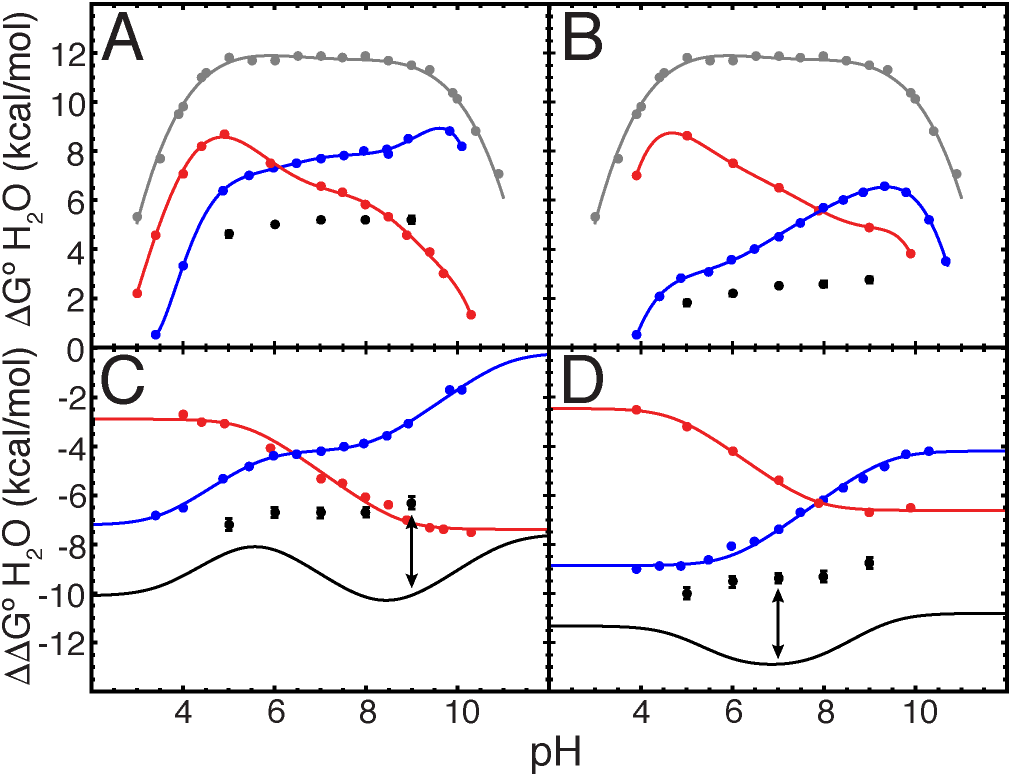
pH dependence of the thermodynamic stabilities of the KE (**A and C**) and EK (**B and D**) variants. Data for single Lys variants are shown in blue, single Glu variants in red, double variants in black, and background protein in grey. Solid lines in A and B are meant solely to guide the eye. Solid colored lines in C and D show fits to a thermodynamic linkage equation used to determine the p*K*_a_ values of single variants, while black line represents an additive model where there is no coupling energy between the two introduced residues.

The ΔG°_H2O_ of the KE variant ranged from 4.6 to 5.2 kcal/mol at pH values between 5 to 9, with a stability of 5.2 kcal/mol at pH 7. The ΔG°_H2O_ of the EK variant ranged from 1.8 to 2.7 kcal/mol at pH values between 5 to 9, with a stability of 2.5 kcal/mol at pH 7. The thermodynamic stabilities of the KE and EK variants relative to the background protein (ΔΔG°_H2O_) are relatively constant between pH 5-9. These data are consistent with the p*K*_a_ value of both groups being either normalized or further shifted in opposite directions (equal but opposite p*K*_a_ shifts would destabilize the protein by equal but opposite amounts, canceling out the pH-dependence of the protein’s thermodynamic stability^12^).

The double variants were significantly more stable than expected from the introduction of two destabilizing substitutions. The coupling energy (ΔΔG_int_) was determined by comparing the relative stabilities of these variants compared to background to an additive model of the single variants, which assumes no interaction between the two residues^23^:

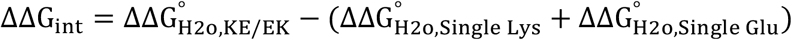

At the pH values where the additive model is predicted to be most destabilizing (~pH 9 for KE and ~pH 7 for EK), the KE and EK variants are 3.9 and 3.4 kcal/mol more stable than predicted, respectively. It is important to note that these coupling energies do not reflect a direct interaction between the two residues, as the assumption that the protein structure is not perturbed by introduction of the two ionizable side chains was shown to not be correct (see below).

### Crystal structures

Crystal structures of the KE and EK variants were solved to 1.7 and 2.2 Å, respectively, allowing for a structural comparison with the background protein and single variants (Figure 3A-C). The microenvironment of Thr-62 and Val-66 is completely hydrophobic in the background protein (Figure 3D). In the T62K variant, Lys-62 is completely buried and excluded from solvent (Figure 3E), while the ionizable moieties of the buried residues in T62E, V66E, and V66K are within hydrogen bonding distance (<3.5 Å) of buried or interfacial water molecules (Figure 3F-H).

**Figure 3.**
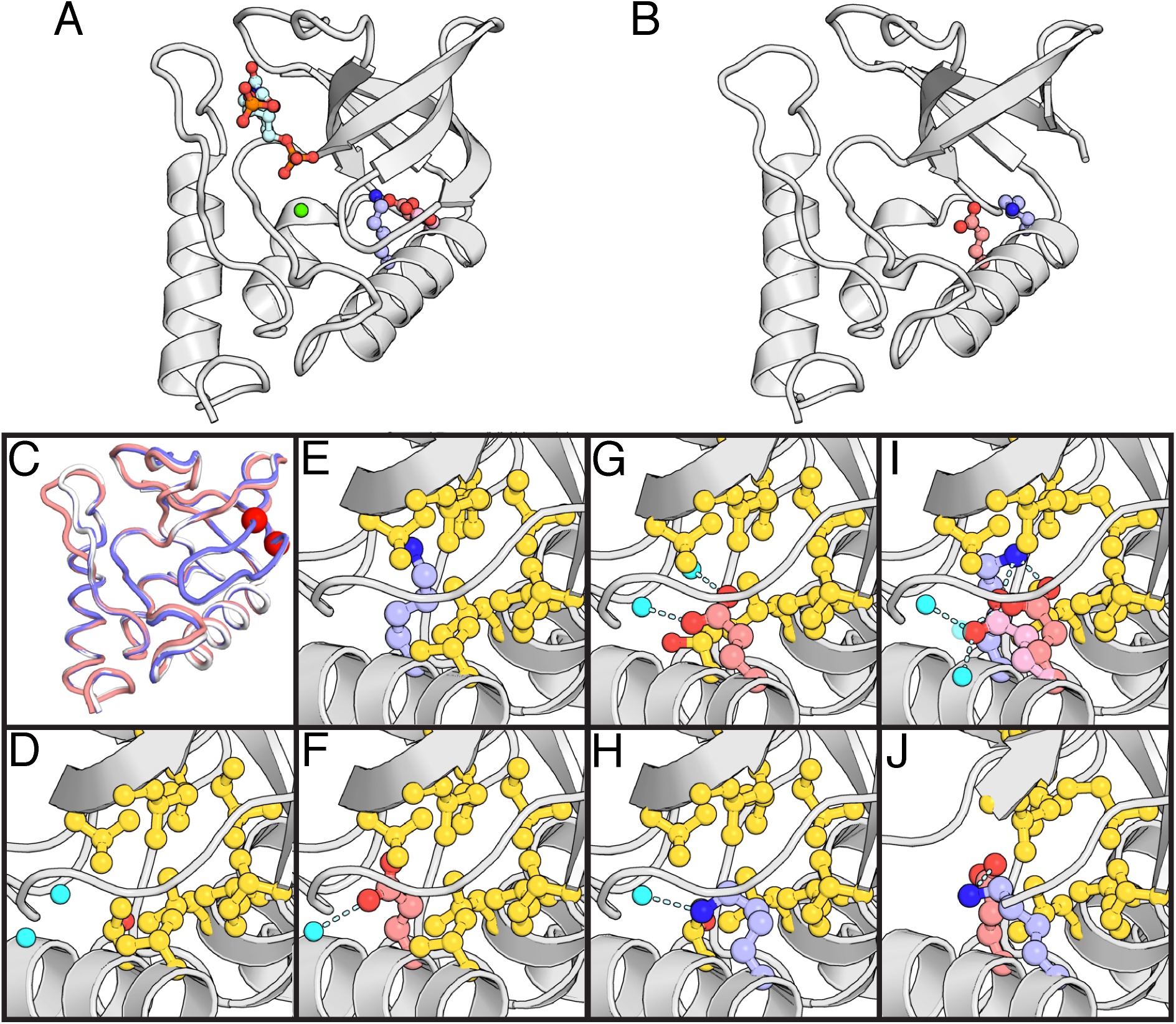
Crystal structures. (**A**) Crystal structure of KE at pH 9, in complex with calcium and inhibitor pdTp. (**B**) Crystal structure of EK at pH 7. (**C**) Backbone overlay of background protein at pH 8 in white, KE in light blue, and EK in salmon. Region of EK crystal structure where electron density is missing is highlighted with red spheres. (**D-J**) Microenvironment of 62:66 pairs in the background protein at pH 8 (**D**), T62K at pH 8 (**E**), V66E at pH 6 (**F**), T62E at pH 9 (**G**), V66K at pH 9 (**H**), KE at pH 9 (**I**), and EK at pH 7 (**J**). Hydrophobic residues are highlighted in yellow, and water molecules in cyan. Transparent water molecule in KE is partial occupancy. Lower occupancy Glu-66 side chain in KE is shown in light pink.

The crystal structure of KE was solved in complex with Ca^2+^ and thymidine 3’,5’- diphosphate (pdTp) at pH 9 (Figure 3A, I). The protein backbone is superimposable with that of the background protein (Figure 3C). Lys-62 and Glu-66 are buried and within hydrogen bonding distance to each other, though Glu-66 is observed in two conformations. In the more deeply buried conformation (55% occupancy) the Oε1 and Oε2 atoms of Glu-66 are 2.6 and 3.4 Å away from the Nζ atom of Lys-62, and a single internal water molecule is found 2.7 Å away Oε2. The minor conformation of Glu-66 (45% occupancy) is more solvent exposed with Oε2 overlapping the position of the H-bonded water in the major conformation; Oε1 is within 2.8 Å from the Nζ atom of Lys-62, while Oε2 is 2.7 and 3.1 Å from two interfacial water molecules. Lys-62 makes no other polar contacts besides those to Glu-66.

The crystal structure of EK was solved at pH 7 in the absence of inhibitor and showed evidence of significant deviation from the background protein (Figure 3B, J). This variant crystallized in a P6(3) space group, which has not been observed in any of the 250+ previously solved structures of SNase. The loop region between residues 113-120 adopts a unique conformation due to different intermolecular contacts in this crystal form. Electron density for residues 15-24 in the β-1 and β-2 strands is very weak or missing and could not be reliably modeled. This structural change effectively exposes Glu-62 and Lys-66 to bulk solvent. The ionizable moieties of Glu-62 and Lys-66 are shifted slightly relative to their positions in the single variants to be in hydrogen bonding distance from each other, with an Oε to Nζ distance of 3.2 Å.

### NMR spectroscopy

#### Backbone characterization

The ^1^H-^15^N HSQC spectra of both KE and EK variants contain well-dispersed, sharp resonances, indicative of folded proteins, though a small but significant degree of random coil resonances were observed in the EK spectra (Figure 4A, B). Each variant had a single resonance missing that was assignable in the background protein (residues 23 and 21 in EK and KE, respectively). Chemical shift perturbations (CSPs) were observed for both the backbone N and H_N_ resonances with the largest changes occurring around the mutation site and the β-1 and β-2 strands, the latter region being where resonance loss was observed (Figure S2-3). Similar trends were observed for the Cα CSPs, which are highly sensitive to structural perturbations in proteins^24^, in both variants near the β-1 and β-2 strands (Figure 4C, D, S4). The CSPs around the β-1 and β-2 strands in all nuclei were larger in the EK variant. ^1^H-^15^N NOE measurements demonstrate that the β-1/β-2 turn was more dynamic on the ps-ns timescales in EK than in KE or the background protein, consistent with partial unfolding (Figure 4E). The chemical shifts of the residues in this turn in EK are consistent with partial unfolding, based on Random Coil Index predictions (Figure 4F).

**Figure 4.**
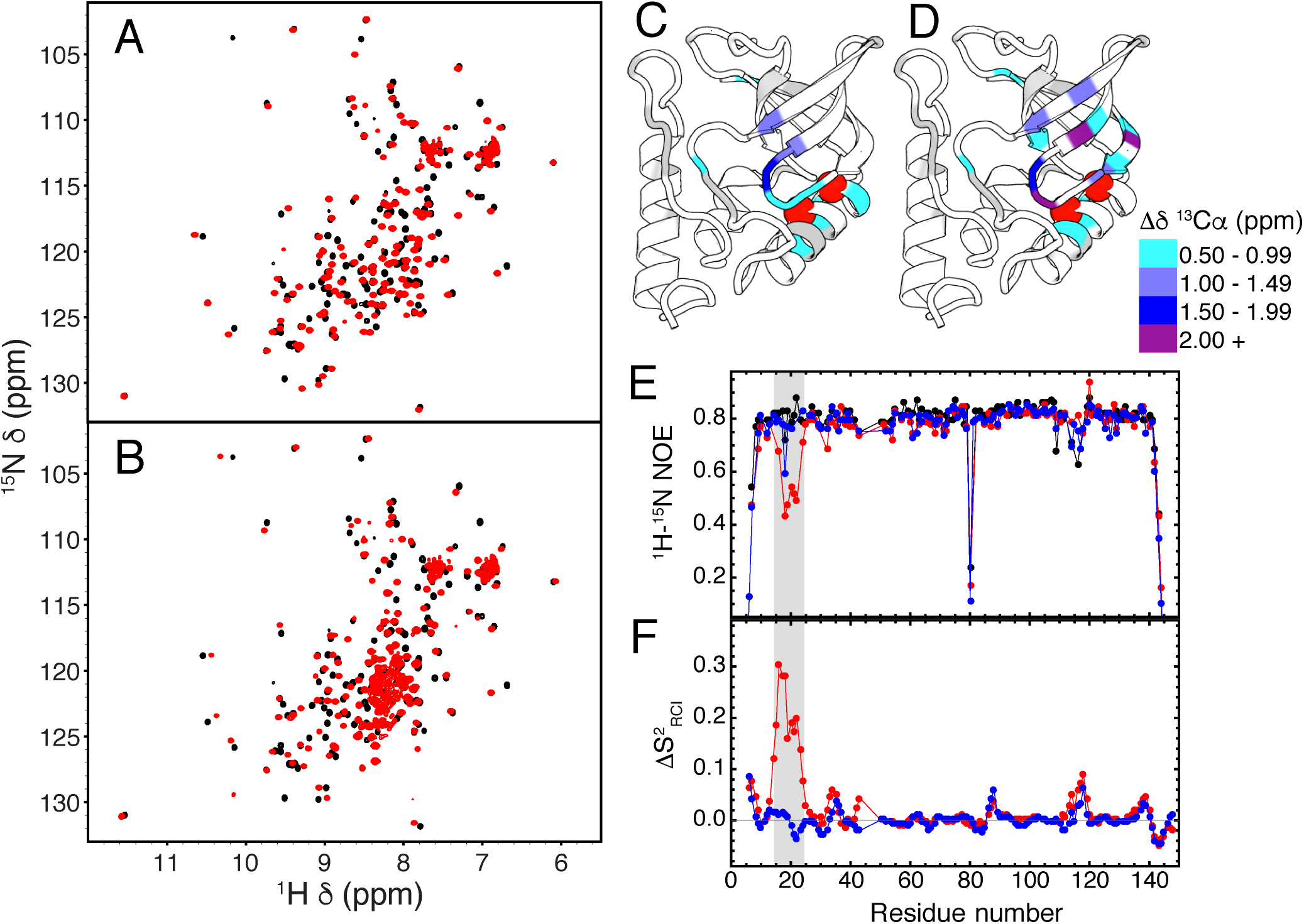
Backbone characterization by NMR spectroscopy. ^1^H-^15^N HSQC spectra of background protein (pH 6.6), (**A**) KE (pH 7.0), and (**B**) EK (pH 6.6) variants. The pH value of each sample is denoted in parenthesis. ^13^Cα chemical shift perturbations (CSPs) of (**C**) KE, pH 7.0 and (**D**) EK, pH 6.6, relative to the background protein at pH 6.6. The mean CSP was 0.21 for KE and 0.28 for EK, and the standard deviation was 0.48 for KE and 0.60 for EK. Substitution sites are shown as red spheres, but are not included in analysis due to the sensitivity of the Cα chemical shift to the residue identity. (E) ^1^H-^15^N heteronuclear NOE analysis of background protein (pH 7.4, black), KE (pH 6.1, blue), and EK (pH 7.4, red). The pH values were chosen based on spectral resolution. (F) Comparison of Random Coil Index S^2^ values (predicted from chemical shifts), shown as background (pH 6.6) minus variant (ΔS^2^). ΔS^2^ values for KE variant (pH 7.0) are shown in blue, and those for EK (pH 6.6) in red. More positive values indicate regions that are more dynamic in variant than the background protein. Values below 0.8 should not be considered significant.

#### Direct detection of buried ion pairs with NMR

Side chain Cβ/γ-CO correlation spectra (^13^C-^13^C CBCGCO) of the single Glu variants and ion pair variants (Figure 5A, B, S5) were collected to investigate the charge state and hydrogen bonding patterns of the introduced side chain carboxylate moieties. The chemical shifts of the neutral Glu resonances in the single variants were distinct from those of the surface residues. The chemical shifts of E62 and E66 in the double variants were more similar to those of surface residues than the buried, neutral Glu residues, and very similar to the previously studied ionized E23 in the V23E/L36K variant of SNase^12^. Two resonances in the CBCGCO spectra of KE were assignable to Glu-66, indicative of slow exchange behavior caused by conformational exchange on ms or longer timescales^25^ consistent with the alternate conformations observed in the crystal struture. In the double variants the ^13^C chemical shift of the side chain carboxylate of both E62 and the major form of E66 did not show evidence of a deuterium isotope shift upon transfer from H_2_O to D_2_O, demonstrating that these Glu residues are deprotonated between pH 7-8. The resonance of the minor form of E66 was too broad for isotope shift studies. Both the major and minor form of E66 and E62 experience intermediate to slow exchange below pH 6, presumably with the protonated state.

**Figure 5.**
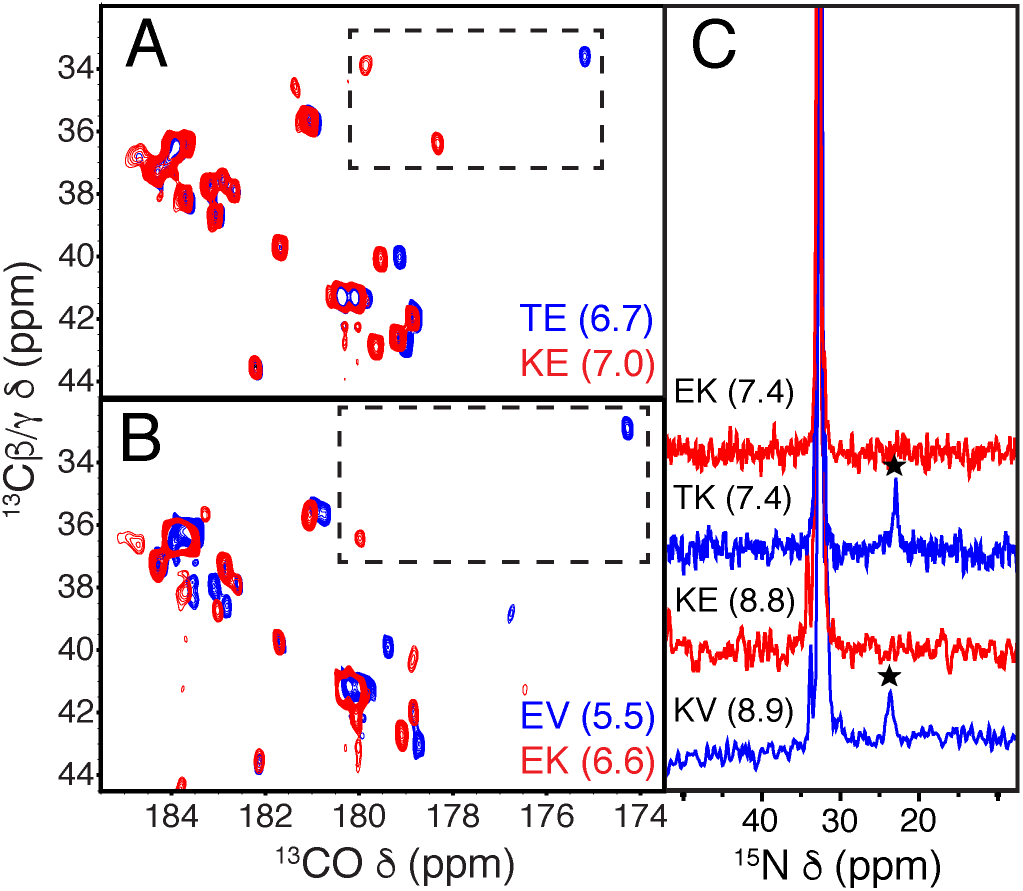
NMR detection of buried ion pairs. ^13^C-detect CBCGCO spectra of (**A**) KE and (**B**) EK single and double variants. Resonances from E66/E62 are highlighted in dashed boxes. The pH for each spectra were chosen based on spectral quality; changes in the position of E62 in single variant (EV) between pH 5.5 and 6.6 were minimal (Figure S6). (**C**) ^15^N 1D spectra of T62K (KV) and KE (pH 8.9, bottom) and V66K (TK) and EK (pH 7.4, top). Resonances belonging to deprotonated Lys residues are denoted with stars, while the protonated Lys and the N-terminal NH_3_+ groups resonate between 30-40 ppm.

The protonation state of the introduced Lys residues was determined by detection of their side chain ^15^N and ^13^C nuclei. At pH values where Lys-62 and Lys-66 are deprotonated in the single variants, no Nζ resonances consistent with a deprotonated Lys (20-26 ppm^26,27^) residue are observed in the double variants in ^15^N 1D experiments (Figure 5C), suggesting that either the Lys residues are protonated and indistinguishable from the surface resonances, or are broadened beyond detection due to μs-ms dynamics. To distinguish the between those possibilities, the side chain carbon resonances were tracked as well, as the ^13^Cδ chemical shift is also a sensitive reporter of charge state^1^. All side chain ^13^C resonances of Lys-62 were broadened beyond detection past the ß position, suggestive of dynamic behavior. Similar behavior is observed previously in the T62K single variant when Lys-62 ionizes^27^ (Figure S7). In contrast, the Cδ and Cε resonances of Lys-66 were observed in CC(CO)NH and ^1^H-^13^C-^13^C NOESY experiments. The Cδ of Lys-66 was shifted upfield by 4.5 ppm in the EK variant relative to its value in the V66K variant at pH values where Lys-66 should be neutral. The magnitude of the shift is consistent with the ionization of the Lys side chain^28,29^. NOEs from deeply buried methyl groups to Lys-66 were also observed in both the V66K and EK variants, consistent with groups being buried in the hydrophobic protein interior as observed in the crystal structures (Figure S8).

## Discussion

Classic continuum electrostatic theory describes the energetics of ion pair burial through the self-energies experienced by dehydrating each charge (ΔG_ii_) and the Coulomb interaction experienced by the two charges (ΔG_ij_). These energies are sensitive to the dielectric constant of the surrounding medium; although the measured dielectric constant of dry protein powders is between 2-4^30^, continuum calculations require artificially high dielectric constants to accurately reproduce electrostatic effects in proteins^31^. The origin of the high dielectric constants has been controversial, having been attributed to factors including water penetration^32,33,34^, side chain heterogeneity^35,36^, and backbone reorganization^37,38,27^. Previous structural and thermodynamic characterization two ion pairs in SNase^12,14^, along with the two additional pairs studied here has demonstrated that backbone reorganization is necessary to accommodate these groups in their charged state. The novel insight this study provides is the structural detail into how proteins reorganize to accommodate buried charges; the range of conformational responses observed in this study illustrates the challenges that structure-based energy calculations must overcome to properly capture the complexities of the local structural responses in response to charge burial.

## Dielectric breakdown of the hydrophobic core

The EK and KE variants in this study were observed to undergo reorganization by CD, NMR spectroscopy and X-ray crystallography. The crystal structure and NMR spectroscopy data are completely consistent with partial unfolding of the β-1/β-2 region of the EK variant, which exposes the buried residues to bulk solvent. In the KE variant, the crystal structure showed both residues are internal, with NMR evidence suggesting only minor reorganization of the ß-1/ß-2 region. Instead, conformational heterogeneity of the Glu-66 side chains was observed in KE structure and supported by intermediate and slow exchange behavior of side chain by NMR. This suggests side chain dynamics allow the residues to sample well-hydrated interfacial positions. Although it has been previously demonstrated that Lys residues are capable of rapidly making and breaking hydrogen bonds on ps-ns timescales^39,40,41^, the Lys:Glu pair KE appear to exhibit exchange at much slower timescales (μs-s). The slow exchange may reflect a large energy barrier between conformations due to the presence of dehydrated intermediate states. Finally NMR spectroscopy and analysis of the pH-dependence of the thermodynamic stability of these variants demonstrates that the Lys and Glu residues are charged between pH 5-9 suggesting the observed structural change is sufficient to shift the p*K*_a_ values of the groups away from their values in the single variants and back towards their values in bulk solvent.

The degree of reorganization observed in the EK and KE variants show that the dielectric environment governing the interaction between the charged residues is not governed by the properties of the protein interior, but rather the protein-water interface. In this sense, the responses of the EK and KE variants both represent dielectric breakdown as the protein interior must reorganize to the extent that its properties no longer dictate the interaction of the charged residues^42^. Instead, the localized structural responses in both variants are necessary to effectively hydrate the buried residues, allowing the residues to behave as if they were in water.

## The conformational ensemble modulates electrostatic interactions in the protein interior

It is important to note that the ion pairs investigated here and in previous studies all perturb the same structural region of the protein, the β-1/β-2 strands and the intervening hairpin turn, though the magnitude of reorganization varies dramatically between the four variants. Crystal structures and NMR spectroscopy have shown that the β-1/β-2 strands can undergo partial unfolding in response to the ionization of a single Glu residue^38^, although more subtle responses localized to just the hairpin turn between the strands have been observed^27^ (Figure 6A-B). In variants where E23/K36 or K62/E66 pairs are introduced the backbone response is subtle, with similar chemical shift perturbations in both variants localized to the β-1/β-2 turn (Figure 6C-D, S4). Subtle backbone reorganization around this β-turn in these double variants may be underestimated in crystal structures. Both structures are bound to inhibitor and calcium, the latter requiring residues on the β1/2-turn for binding. Although the backbone behavior of variants with E23/K36 and K62/E66 are similar, the side chains in the protein interior have very different behaviors. The E23/K36 pair is stabilized by a “polar cage” of internal water molecules and protein dipoles. In the K62/E66 pair, both side chains show evidence of conformational heterogeneity; minor backbone reorganization alone cannot effectively stabilize the internal pair, the side chains must sample multiple conformations to remain effectively hydrated.

**Figure 6.**
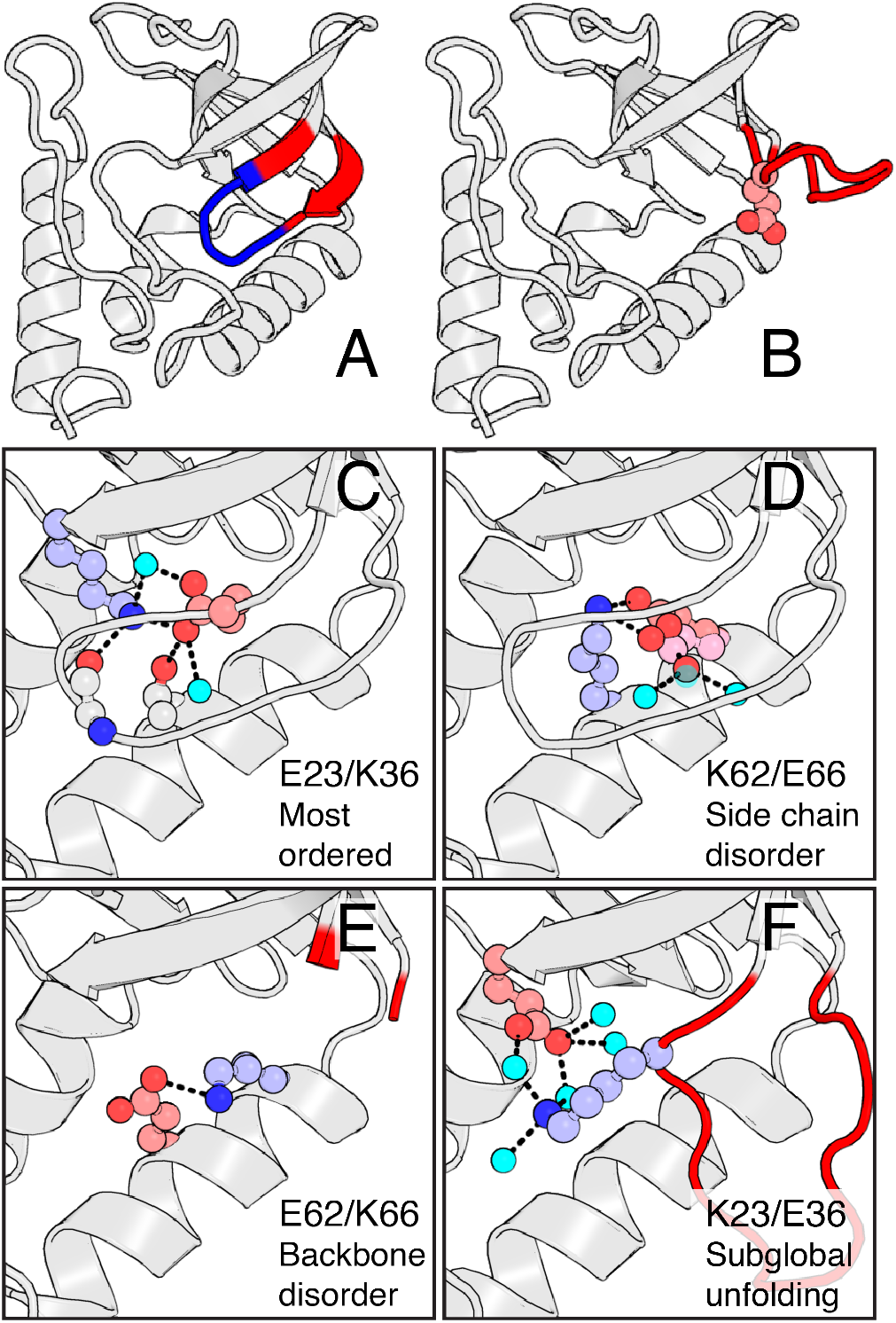
Conformational modulation of electrostatic energies in the protein interior. (**A**) Regions of β-1/β-2 showing evidence of reorganization (by ^13^Cα CSP) in variants with E23/K36 and K62/E66 (blue), and in variant with E62/K66 (red). (**B**) Crystal structure of variant with E23 at pH values where E23 is ionized.^38^ (**C-F**) Crystal structures of the four ion pair variants demonstrating range of conformational responses, from least to most structured variants. The structure of the variant with K23/E36 was solved in a more stable background of SNase.

These subtle responses are in stark contrast to the backbone response to the ionization of E62/K66, which leads to unfolding of a 10 residue stretch of the β-1/β-2 strands, and K23/E36, which triggers subglobal unfolding (Figure 6E-F). The K23/E36 pair must be introduced into a more stable variant of SNase, in order to be folded in solution. Even then, the protein responds with a similar structure response as observed in the variant with E62/K66, where a portion of β-1/β-2 is unfolded but the rest of the protein is unperturbed^14^. In this state, K23 and E36 do not form a direct, “contact” ion pair^41^ in the crystal structure, but appear to instead form a solvent-separated ion pair^43^.

Although each variant studied has a different mechanism for tolerating burial of the charged pairs, in every case some degree of backbone reorganization was necessary, either as a dielectric response – making the protein interior a better solvent, or as dielectric breakdown – reorganization to expose the buried residues to solvent. What is remarkable is that these different behaviors are modulated by the same structural element in the protein, which can adopt multiple conformations to stabilize the ion pairs. The inherent conformational heterogeneity of this region of the protein is likely to be important for the success of these ion pair studies in SNase. Not unsurprisingly, the energetics of ion pair burial appear to be more complex than of burial of single ionizable groups. In a survey of 25 internal Lys substitution in SNase, the cost of ionizing the buried side chain was correlated to the cost of ionizing the buried Lys and the extent of structural reorganization (manuscript in preparation). In contrast, the most ordered protein in the ion pair studies, the E23/K36 variant, is as stable as the E62/K66 variant (ΔG of both proteins = 2.5 kcal/mol at pH 7), despite the latter protein undergoing significant partial backbone unfolding. On the other hand, the K62/E66 variant is the most stable of the four variants studied (ΔG = 5.2 kcal/mol at pH 7). yet shows evidence of significant side chain and minor backbone reorganization.

## Implications

The results of this study demonstrate that conformational heterogeneity can and does modulate charge-charge interactions in the protein interior. Previous statistical and structure-based calculation studies have suggested that buried ion pairs in nature can be stabilized through Coulomb interactions^44,45^. At least in SNase, Coulomb interactions alone are insufficient to stabilize the ionization of buried residues; the protein interior must either reorganize to increase its polarity or the buried residues must be exposed to bulk solvent. Both processes involve some degree of structural reorganization. The results of this study contradict a recent computational study that suggests ionized acidic residues in the protein interior can be stabilized by pairing with hydronium ions^46^. Although methods that explicitly account for reorganization have already been used to study the backbone response of the E23/K36 variant, it represents perhaps the simplest case given the relatively minor magnitude of the response ^47,48^. The highly local, yet varied nature of the backbone and sidechain conformational responses of the variants in this study will be difficult to capture using current structure-based methods for studying pH-dependent conformational changes in proteins^49,50,51,52,53^.

In nature, dynamical processes involving ion pairs in interfacial and hydrophobic environments are essential for a wide range of biochemical processes including both H^+^ transport and e^-^ transfer and for modulating protein dynamics and pH-sensitivity in proteins. Ion pairs in protein-protein interfaces have been shown to be important for driving allosteric responses ^54,55^. Enzyme active sites also utilize buried ion pairs to modulate the reactivity of nucleophiles, tune the p*K*_a_ values of general acids/bases, and stabilize transition states. Although dynamics have been invoked to explain the catalytic power of some enzymes^56,57,58,59^, the physical basis for how these motions are related to catalysis remains controversial^60^. Nevertheless, experimental work has suggested that evolution of effective enzymes requires a balance of stability and flexibility to effectively tune function^61^ for specific environmental niches^62^. The results of this study suggest that conformational heterogeneity or flexibility can be used as a mechanism for tuning electrostatic energies for their functional roles. Although the timescales of motions in this study are too slow to be relevant to catalysis, the relationship between flexibility^63,61^ and stability^62^, and the propensity of proteins to sample alternative states in response to charge burial, must be accounted for in the de novo design of novel enzymes.

## Conclusions

The role of conformational reorganization in modulating the dielectric properties of proteins is of great interest in understanding biological energy transduction, catalysis, and pH-sensing. The results of this study indicate the challenges facing structure based energy calculations and design algorithms; although it may be possible to design proteins with enough thermostability to tolerate the introduction of functional sites, the arrangement of an active site in a crystal structure or the determination of the lowest energy conformation in structure prediction algorithm will not accurately capture the range of conformations observed in solution. Rather, methodologies that seek to explicitly model the conformational heterogeneity of proteins will likely prove to be more successful in the design of novel proteins with the ability to drive energy transduction, respond to changes in pH, and catalyze novel chemical reactions. This will be difficult, as the highly local nature of the structural responses observed in this study, which range from side chain disorder to backbone unfolding, will be difficult to accurately capture for even the most state of the art structure-based energy calculation methods.

## Materials and Methods

### Protein production and purification

Introduction of the T62E/V66K and T62K/V66E substitutions into the hyperstable A+PHS variant of SNase was performed as previously described with primers purchased from Integrated DNA Technologies (IDT)^1^. Unlabeled and ^15^N or ^13^C/^15^N labeled proteins were purified as previously described, using ^15^N ammonium chloride or ^13^C glucose purchased from Cambridge Isotope Labs^64^.

### Circular dichroism and fluorescence spectroscopy

Protein samples consisted of 100 mM KCl, 10 mM buffer (CD) or 25 mM buffer (fluorescence), and 125 μg/ml (CD) or 50 μg/ml (fluorescence) protein. Potassium acetate was used as a buffer for pH 5 measurements, MES for pH 6, HEPES for pH 7 (substituted with Tris for CD due to high background signal of HEPES), TAPS for pH 8, and CHES for pH 9. All CD experiments were performed on an Aviv model 420 circular dichroism spectrometer, as previously described, except each point (collected every 1 nm between 195-300 nm) was averaged for 10 seconds. All guanidinium melts were performed on an Aviv model 107 Automated Titration Fluorimeter as previously described, using ultra-pure guanidinium hydrochloride purchased from Fischer. All data was collected at 25 °C.

### NMR Spectroscopy

Assignment and ^13^C detected experiments were performed on Bruker Avance or Avance II spectrometers operating at a 600 MHz ^1^H frequency, equipped with TCI cryogenic probes. NOESY experiments were performed on a Varian 800 MHz Inova spectrometer with a room temperature probe. ^15^N detected data were collected on Varian 500 MHz Inova or Bruker 400 MHz Avance III spectrometers, both with room temperature broadband observe probes. Protein samples consisted of 0.9-1.2 mM protein in 25 mM buffer (HEPES pH 7.0 for KE, MES pH 6.6 for EK), 100 mM KCl, and 10% D2O for assignments. For ^1^H-^15^N heteronuclear NOE experiments, data was collected at pH 6.1 in 25 mM MES for the KE variant, and pH 7.4 in 25 mM HEPES for the EK variant. The pH 7.4 sample of EK was also used for NOESY experiments. Assignments were obtained using standard triple resonance experiments (HNCACB, CBCA(CO)NH, HNCO, HBHA(CO)NH, H(CCCO)NH, CC(CO)NH). A CCH-TOCSY was necessary to assign the minor Glu-66 resonance in KE. NOESY experiments were performed using CH3 selective ^1^H-^13^C-^13^C NOESY-HMQC and ^1^H-^13^C-^1^H NOESY-HSQC pulse sequences. ^1^H-^15^N NOE experiments were performed with a 4 second saturation period or recycle delay. NOE was calculated as the ratio of resonance intensities with versus without NOE saturation. ^13^C detect CBCGCO experiments were performed using a previously published pulse sequence^38^. 64-128 scans were necessary to detect the resonances of E62 and E66, with acquisition times of 20 ms in the indirect dimension. For deuterium isotope shift experiments, a sample of each variant was made in 25 mM HEPES and 100 mM KCl (pH 7.3 and 7.8 for KE and EK, respectively). Samples were split into two tubes of identical volume (400 ml), lyophilized, and then resuspended in equivalent volumes of either 90% H_2_O/10% D_2_O or 99% D_2_O. ^15^N 1D and heteronuclear cross polarization experiments were performed using the same parameters as previously described^27^. 1D data was visualized using Topspin 3.7. 2D and 3D data were processed with the NMRpipe suite, and analyzed using Sparky 3.115. All data was collected at 25 °C.

### Crystallography

Crystals of T62E, V66E, and KE (all in the Δ+PHS background) were grown using the hanging-drop vapor diffusion method at 4 °C. Protein concentrations of 8-12 mg/ml were used for each protein. Crystals of EK were grown at room temperature (20 °C) with a concentration of 23 mg/ml. 4 μl of protein solution was mixed with 4 μl of mother liquor. Crystals appeared within a month, and all were flash frozen prior to data collection except for the EK crystals, which were pushed through paratone oil prior to freezing and data collection. Details of data collection and statistics are provided in Table Sxx. All data was indexed, integrated, scaled and merged using the manufacturer’s software. Processing was performed using the CCP4 suite, with initial phasing by molecular replacement done using PHASER, with the background protein’s crystal structure used for the starting model (PDB ID: 3BDC). Model building and refinement was performed using COOT and Refmac-5, respectively.

## Supporting Information

Tables of thermodynamic parameters from chemical denaturation experiments (S1-3) and crystallographic statistics (S4-5). Circular dichroism spectra of single variants, backbone chemical shift perturbations, deuterium isotope shift and pH dependent behaviors of Glu side chains by CBCGCO, and detection of Lys carbon resonances (S1-8).

## Acknowledgements

The authors would like to thank Dr. Katherine Tripp for assistance with CD data collection, Dr. Maxime Siegler for assistance with crystallography data collection, and Dr. Aaron Robinson and Mr. Matthew Sternke for initial NMR studies. Data was collected at the Johns Hopkins University Biomolecular NMR Center (500, 600 and 800 MHz data), Department of Chemistry NMR Facility (400 MHz data), Center for Molecular Biophysics (circular dichroism experiments), and Department of Chemistry X-Ray Crystallography Facility. Funding for this work was provided by the NIH (grant GM 061597 to BGME) and NSF (grant MCB1517378 to BGME).

## References

1. Isom, D. G., Castañeda, C. A., Cannon, B. R., Velu, P. D. & E, B. G.-M. Charges in the hydrophobic interior of proteins. Proc. Natl. Acad. Sci. 107, 16096–16100 (2010).

2. Harris, T. K. & Turner, G. J. Structural Basis of Perturbed pKa Values of Catalytic Groups in Enzyme Active Sites. IUBMB Life 53, 85–98 (2002).

3. Schwans, J. P., Sunden, F., Gonzalez, A., Tsai, Y. & Herschlag, D. Uncovering the Determinants of a Highly Perturbed Tyrosine pKa in the Active Site of Ketosteroid Isomerase. Biochemistry 52, 7840–7855 (2013).

4. Lanyi, J. K. Proton transfers in the bacteriorhodopsin photocycle. Biochim. Biophys. Acta BBA - Bioenerg. 1757, 1012–1018 (2006).

5. Ballmoos, C. von, Wiedenmann, A. & Dimroth, P. Essentials for ATP Synthesis by F1F0 ATP Synthases. Annu. Rev. Biochem. 78, 649–672 (2009).

6. Hong, H., Szabo, G. & Tamm, L. K. Electrostatic couplings in OmpA ion-channel gating suggest a mechanism for pore opening. Nat. Chem. Biol. 2, 627 (2006).

7. Isom, D. G. et al. Protons as Second Messenger Regulators of G Protein Signaling. Mol. Cell 51, 531–538 (2013).

8. Isom, D. G., Sridharan, V. & Dohlman, H. G. Regulation of Ras Paralog Thermostability by Networks of Buried Ionizable Groups. Biochemistry 55, 534–542 (2016).

9. Barlow, D. J. & Thornton, J. M. Ion-pairs in proteins. J. Mol. Biol. 168, 867–885 (1983).

10. Rashin, A. A. & Honig, B. On the environment of ionizable groups in globular proteins. J. Mol. Biol. 173, 515–521 (1984).

11. Hendsch, Z. S. & Tidor, B. Do salt bridges stabilize proteins? A continuum electrostatic analysis. Protein Sci. 3, 211–226 (1994).

12. Robinson, A. C., Castañeda, C. A., Schlessman, J. L. & E, B. G.-M. Structural and thermodynamic consequences of burial of an artificial ion pair in the hydrophobic interior of a protein. Proc. Natl. Acad. Sci. 111, 11685–11690 (2014).

13. Hwang, J.-K. & Warshel, A. Why ion pair reversal by protein engineering is unlikely to succeed. Nature 334, 270 (1988).

14. Robinson, A. C., Schlessman, J. L. & García-Moreno E, B. Dielectric Properties of a Protein Probed by Reversal of a Buried Ion Pair. J. Phys. Chem. B 122, 2516–2524 (2018).

15. Marqusee, S. & Baldwin, R. L. Helix stabilization by Glu-…Lys+ salt bridges in short peptides of de novo design. Proc. Natl. Acad. Sci. 84, 8898–8902 (1987).

16. Lyu, P. C., Marky, L. A. & Kallenbach, N. R. The role of ion pairs in .alpha. helix stability: two new designed helical peptides. J. Am. Chem. Soc. 111, 2733–2734 (1989).

17. Scholtz, J. M., Qian, H., Robbins, V. H. & Baldwin, R. L. The energetics of ion-pair and hydrogen-bonding interactions in a helical peptide. Biochemistry 32, 9668–9676 (1993).

18. Sivaramakrishnan, S., Spink, B. J., Sim, A. Y. L., Doniach, S. & Spudich, J. A. Dynamic charge interactions create surprising rigidity in the ER/K α-helical protein motif. Proc. Natl. Acad. Sci. 105, 13356–13361 (2008).

19. Baker, E. G. et al. Local and macroscopic electrostatic interactions in single α-helices. Nat. Chem. Biol. 11, 221 (2015).

20. Meuzelaar, H., Vreede, J. & Woutersen, S. Influence of Glu/Arg, Asp/Arg, and Glu/Lys Salt Bridges on α-Helical Stability and Folding Kinetics. Biophys. J. 110, 2328–2341 (2016).

21. Fossat, M. J. & Pappu, R. V. q-Canonical Monte Carlo Sampling for Modeling the Linkage between Charge Regulation and Conformational Equilibria of Peptides. J. Phys. Chem. B 123, 6952–6967 (2019).

22. Isom, D. G., Castañeda, C. A., Cannon, B. R., and E, B. G.-M. (2011) Large shifts in pKa values of lysine residues buried inside a protein. Proc. Natl. Acad. Sci. 108, 5260–5265.

23. Schreiber, G. & Fersht, A. R. Energetics of protein-protein interactions: Analysis ofthe Barnase-Barstar interface by single mutations and double mutant cycles. J. Mol. Biol. 248, 478–486 (1995).

24. Williamson, M. P. Using chemical shift perturbation to characterise ligand binding. Prog. Nucl. Magn. Reson. Spectrosc. 73, 1–16 (2013).

25. Kleckner, I. R. & Foster, M. P. An introduction to NMR-based approaches for measuring protein dynamics. Biochim. Biophys. Acta BBA - Proteins Proteomics 1814, 942–968 (2011).

26. Takayama, Y., Castañeda, C. A., Chimenti, M., García-Moreno, B. & Iwahara, J. Direct Evidence for Deprotonation of a Lysine Side Chain Buried in the Hydrophobic Core of a Protein. J. Am. Chem. Soc. 130, 6714–6715 (2008).

27. Kougentakis, C. M. et al. Anomalous Properties of Lys Residues Buried in the Hydrophobic Interior of a Protein Revealed with 15N-Detect NMR Spectroscopy. J. Phys. Chem. Lett. 9, 383–387 (2018).

28. Platzer, G., Okon, M. & McIntosh, L. P. pH-dependent random coil 1H, 13C, and 15N chemical shifts of the ionizable amino acids: a guide for protein pKa measurements. J. Biomol. NMR 60, 109–129 (2014).

29. Hass, M. A. S. & Mulder, F. A. A. Contemporary NMR Studies of Protein Electrostatics. Annu. Rev. Biophys. 44, 53–75 (2015).

30. Harvey, S. C. & Hoekstra, P. Dielectric relaxation spectra of water adsorbed on lysozyme. J. Phys. Chem. 76, 2987–2994 (1972).

31. Schutz, C. N. & Warshel, A. What are the dielectric “constants” of proteins and how to validate electrostatic models? Proteins Struct. Funct. Bioinforma. 44, 400–417 (2001).

32. Damjanović, A., Brooks, B. R. & García-Moreno E., B. Conformational Relaxation and Water Penetration Coupled to Ionization of Internal Groups in Proteins. J. Phys. Chem. A 115, 4042–4053 (2011).

33. Nguyen, D. M., Leila Reynald, R., Gittis, A. G. & Lattman, E. E. X-ray and Thermodynamic Studies of Staphylococcal Nuclease Variants I92E and I92K: Insights into Polarity of the Protein Interior. J. Mol. Biol. 341, 565–574 (2004).

34. Chakrabarty, S. & Warshel, A. Capturing the energetics of water insertion in biological systems: The water flooding approach. Proteins Struct. Funct. Bioinforma. 81, 93–106 (2013).

35. Alexov, E. G. & Gunner, M. R. Incorporating protein conformational flexibility into the calculation of pH-dependent protein properties. Biophys. J. 72, 2075–2093 (1997).

36. Georgescu, R. E., Alexov, E. G. & Gunner, M. R. Combining Conformational Flexibility and Continuum Electrostatics for Calculating pKas in Proteins. Biophys. J. 83, 1731–1748 (2002).

37. Kato, M. & Warshel, A. Using a Charging Coordinate in Studies of Ionization Induced Partial Unfolding. J. Phys. Chem. B 110, 11566–11570 (2006).

38. Robinson, A. C., Majumdar, A., Schlessman, J. L. & García-Moreno E, B. Charges in Hydrophobic Environments: A Strategy for Identifying Alternative States in Proteins. Biochemistry 56, 212–218 (2017).

39. Zandarashvili, L., Li, D.-W., Wang, T., Brüschweiler, R. & Iwahara, J. Signature of Mobile Hydrogen Bonding of Lysine Side Chains from Long-Range 15N–13C Scalar J-Couplings and Computation. J. Am. Chem. Soc. 133, 9192–9195 (2011).

40. Anderson, K. M. et al. Direct Observation of the Ion-Pair Dynamics at a Protein–DNA Interface by NMR Spectroscopy. J. Am. Chem. Soc. 135, 3613–3619 (2013).

41. Chen, C. et al. Dynamic Equilibria of Short-Range Electrostatic Interactions at Molecular Interfaces of Protein–DNA Complexes. J. Phys. Chem. Lett. 6, 2733–2737 (2015).

42. Karp, D. A., Stahley, M. R. & García-Moreno E., B. Conformational Consequences of Ionization of Lys, Asp, and Glu Buried at Position 66 in Staphylococcal Nuclease. Biochemistry 49, 4138–4146 (2010).

43. Yu, B., Pettitt, B. M. & Iwahara, J. Experimental Evidence of Solvent-Separated Ion Pairs as Metastable States in Electrostatic Interactions of Biological Macromolecules. J. Phys. Chem. Lett. 10, 7937–7941 (2019).

44. Kim, J., Mao, J. & Gunner, M. R. Are Acidic and Basic Groups in Buried Proteins Predicted to be Ionized? J. Mol. Biol. 348, 1283–1298 (2005).

45. Bush, J. & Makhatadze, G. I. Statistical analysis of protein structures suggests that buried ionizable residues in proteins are hydrogen bonded or form salt bridges. Proteins Struct. Funct. Bioinforma. 79, 2027–2032 (2011).

46. Wu, X. & Brooks, B. R. Hydronium Ions Accompanying Buried Acidic Residues Lead to High Apparent Dielectric Constants in the Interior of Proteins. J. Phys. Chem. B 122, 6215–6223 (2018).

47. Pathak, A. K. Effect of a buried ion pair in the hydrophobic core of a protein: An insight from constant pH molecular dynamics study. Biopolymers 103, 148–157 (2015).

48. Pathak, A. K. Constant pH molecular dynamics study on the doubly mutated staphylococcal nuclease: capturing the microenvironment. RSC Adv. 5, 94926–94932 (2015).

49. Shi, C., Wallace, J. A. & Shen, J. K. Thermodynamic Coupling of Protonation and Conformational Equilibria in Proteins: Theory and Simulation. Biophys. J. 102, 1590–1597 (2012).

50. Goh, G. B., Laricheva, E. N. & Brooks, C. L. Uncovering pH-Dependent Transient States of Proteins with Buried Ionizable Residues. J. Am. Chem. Soc. 136, 8496–8499 (2014).

51. Zheng, Y. & Cui, Q. Microscopic mechanisms that govern the titration response and pKa values of buried residues in staphylococcal nuclease mutants. Proteins Struct. Funct. Bioinforma. 85, 268–281 (2017).

52. Liu, J., Swails, J., Zhang, J. Z. H., He, X. & Roitberg, A. E. A Coupled Ionization-Conformational Equilibrium Is Required To Understand the Properties of Ionizable Residues in the Hydrophobic Interior of Staphylococcal Nuclease. J. Am. Chem. Soc. 140, 1639–1648 (2018).

53. Damjanovic, A., Miller, B. T., Okur, A. & Brooks, B. R. Reservoir pH replica exchange. J. Chem. Phys. 149, 072321 (2018).

54. Bell, C. E. & Lewis, M. A closer view of the conformation of the Lac repressor bound to operator. Nat. Struct. Mol. Biol. 7, 209 (2000).

55. Zhan, H., Sun, Z. & Matthews, K. S. Functional Impact of Polar and Acidic Substitutions in the Lactose Repressor Hydrophobic Monomer·Monomer Interface with a Buried Lysine. Biochemistry 48, 1305–1314 (2009).

56. Bhabha, G. et al. A Dynamic Knockout Reveals That Conformational Fluctuations Influence the Chemical Step of Enzyme Catalysis. Science 332, 234–238 (2011).

57. Camilloni, C. et al. Cyclophilin A catalyzes proline isomerization by an electrostatic handle mechanism. Proc. Natl. Acad. Sci. 111, 10203–10208 (2014).

58. Hanoian, P., Liu, C. T., Hammes-Schiffer, S. & Benkovic, S. Perspectives on Electrostatics and Conformational Motions in Enzyme Catalysis. Acc. Chem. Res. 48, 482–489 (2015).

59. Kerns, S. J. et al. The energy landscape of adenylate kinase during catalysis. Nat. Struct. Mol. Biol. 22, 124–131 (2015).

60. Warshel, A. & Bora, R. P. Perspective: Defining and quantifying the role of dynamics in enzyme catalysis. J. Chem. Phys. 144, 180901 (2016).

61. Bar-Even, A., Milo, R., Noor, E. & Tawfik, D. S. The Moderately Efficient Enzyme: Futile Encounters and Enzyme Floppiness. Biochemistry 54, 4969–4977 (2015).

62. Sugrue, E., Carr, P. D., Scott, C. & Jackson, C. J. Active Site Desolvation and Thermostability Trade-Offs in the Evolution of Catalytically Diverse Triazine Hydrolases. Biochemistry 55, 6304–6313 (2016).

63. Elias, M., Wieczorek, G., Rosenne, S. & Tawfik, D. S. The universality of enzymatic rate–temperature dependency. Trends Biochem. Sci. 39, 1–7 (2014).

64. Castañeda, C. A. et al. Molecular determinants of the pKa values of Asp and Glu residues in staphylococcal nuclease. Proteins Struct. Funct. Bioinforma. 77, 570–588 (2009).

